# *TeraVR* Empowers Precise Reconstruction of Complete 3-D Neuronal Morphology in the Whole Brain

**DOI:** 10.1101/621011

**Authors:** Yimin Wang, Qi Li, Lijuan Liu, Zhi Zhou, Yun Wang, Lingsheng Kong, Ning Zhong, Renjie Chai, Xiangfeng Luo, Yike Guo, Michael Hawrylycz, Qingming Luo, Zhongze Gu, Wei Xie, Hongkui Zeng, Hanchuan Peng

## Abstract

Neuron morphology is recognized as a key determinant of cell type, yet the quantitative profiling of a mammalian neuron’s complete three-dimensional (3-D) morphology remains arduous when the neuron has complex arborization and long projection. Whole-brain reconstruction of neuron morphology is even more challenging as it involves processing tens of teravoxels of imaging data. Validating such reconstructions is extremely laborious. We developed *TeraVR*, an open-source virtual reality annotation system, to address these challenges. *TeraVR* integrates immersive and collaborative 3-D visualization, interaction, and hierarchical streaming of teravoxel-scale images. Using *TeraVR*, we produced precise 3-D full morphology of long-projecting neurons in whole mouse brains and developed a collaborative workflow for highly accurate neuronal reconstruction.

## Introduction

Major international initiatives are underway to profile and characterize cell types of the mammalian brain (Ecker, 2017, Regev, 2017). As a key recognized attribute of cell type since Ramon y Cajal, high fidelity reconstruction of neuron morphology is gaining increased attention (Ascoli, 2006; Yuste, 2015; Economo, 2016). The basic building blocks of the brain, neurons and glial cells, are often noted for their remarkable three-dimensional (3-D) shapes that distinguish one cell-type from another. While such shapes are critical to understanding cell type, function, connectivity and development (Zeng and Sanes, 2017), it is challenging to profile these shapes precisely. Sparse labeling and high-resolution micro-imaging of a brain cell help visualize the appearance of the cell, yet it remains a major bottleneck how to convert such imaging data into a digital description of morphology, including the 3-D spatial locations of a cell’s parts and their topological connections. This conversion process is often called *neuron tracing* or *neuron reconstruction* and it has become an essential and active area of neuroinformatics.

Two complementary reconstruction workflows exist: one for electron microscopy (EM) images and the other for light microscopy (LM) data (Helmstaedter, 2013; Januszewski, 2018; Peng, et al, 2015). EM offers nanometer-resolution and thus provides a way to reconstruct the entire surface of the shape, but it is often constrained to relatively small brain regions. When whole-brain scale is the focus and complete neuron morphology is desired, LM is a more suitable imaging modality where data is typically acquired at sub-micrometer resolution. LM-reconstruction makes it possible to trace both long projections and the terminal arborization of a brain cell. Recent extension of this approach based on expansion microscopy can help visualize neurons at nanometer-resolution using LM approaches (Gao, 2019).

It is widely recognized that manual and semi-automatic neuron-tracing methods are crucially required to produce full reconstructions, which can also serve as “gold-standard” datasets to develop fully automatic neuron-tracing methods (Peng, 2011; Peng, 2015; Ai-Awami, 2016; Mosinska, 2017; Haehn, 2018). Without loss of generality below we define any neuron-tracing method that has a non-negligible human labor component as *manual reconstruction*, which clearly also includes many semi-automatic methods. This paper discusses a new technology that makes such LM-oriented manual reconstruction more efficient and reliable than existing approaches. This work was motivated by four difficulties detailed below: (1) observability, (2) big data handling, (3) interaction, and (4) validation.

First, a neuron can have a very complex 3-D shape that may contain hundreds or even thousands of fiber-branches especially in dense arbors. Such a high degree of mutual occlusion makes it hard to see how neurite-fibers wire together. The observability is further compromised by the uneven or weak axon labeling, relatively poor Z-resolution from imaging, etc. Often, neither the prevailing 2-D cross-sectional view (such as those widely used in EM-oriented and many LM software packages) nor the typical 3-D intensity projection methods (Peng, et al, 2010) are sufficient to unambiguously delineate these complex wiring patterns, let alone reconstruct them.

Second, reconstructing the full morphology of a mammalian neuron relies on effectively managing and streaming huge whole-brain imaging datasets. The volume of a typical mouse brain is about 500 mm^3^, it is not uncommon that a neuron may have over one hundred millimeters long neurite fiber (Economo, et al, 2016). When an entire mouse brain is imaged at sub-micrometer resolution in 3-D, the volume of the acquired brain images often contains twenty to thirty or more teravoxels. Only a small number of existing software packages are able to open and analyze such big datasets (Bria, et al, 2016; Pietzsch, 2015). How to streamline the unambiguous 3-D visualization and analysis of such huge datasets presents a major informatics challenge.

Third, manual reconstruction of neurons is often laborious and unintuitive using two-dimensional (2-D) tools to interact with 3-D images and the 3-D geometrical representations reconstructed from such images. Reconstructing geometrical objects from 3-D volumetric images requires overlaying these objects onto the imaging data in 3-D space and manipulating them *in situ*. Since most current computer displays (e.g. computer screens) and data interaction tools (e.g. computer mouse) are still restricted to 2-D, it is usually hard to observe and manipulate higher dimensional data via a lower dimensional interface. It is also desirable to interact with the data directly using a smooth workflow. Applications such as *Virtual Finger* (Peng, et al, 2014) represent progress toward this goal, but improvement is still necessary for complex and large neurons and also for display and interaction hardware.

Finally, it is often necessary but very expensive to involve multiple annotators to produce “gold-standard” reconstructions. Manual work is time-consuming and tedious, thus in practice most existing studies can afford only one annotator per neuron. To resolve any ambiguity of reconstructions, it is desired to have a way to allow multiple annotators to visualize the same neuron and its underlying imaging data at the same time, and collaborate on the work. This approach requires collaborative and immersive annotation of multi-dimensional imaging data at the whole-brain scale.

Here we introduce the *TeraVR* system addressing the above requirements. We demonstrate the applicability of *TeraVR* to challenging cases of whole mouse brain neuron reconstruction, achieving previously unattainable accuracy and efficiency.

## Results

We developed *TeraVR* (**Fig. 1, Supplementary Note 1, Supplementary Videos 1-11**), an open-source virtual reality software package for the visualization and annotation of teravoxel-scale whole-brain imaging data (**Fig. 1a**). The software was built upon the *TeraFly* module of *Vaa3D* (http://vaa3d.org) (Bria, et al, 2016), thus *TeraVR* can streamline the data input-output (IO) and other real-time user interaction with teravoxel-scale image volumes, e.g. an 18.4-teravoxel brain-image in **Fig. 1a**. As described below, *TeraVR* also has a number of unique features designed for reconstruction of neuron morphology in whole-brain images, at different levels of details and at different local regions of interest (ROI).

**Figure 1.**
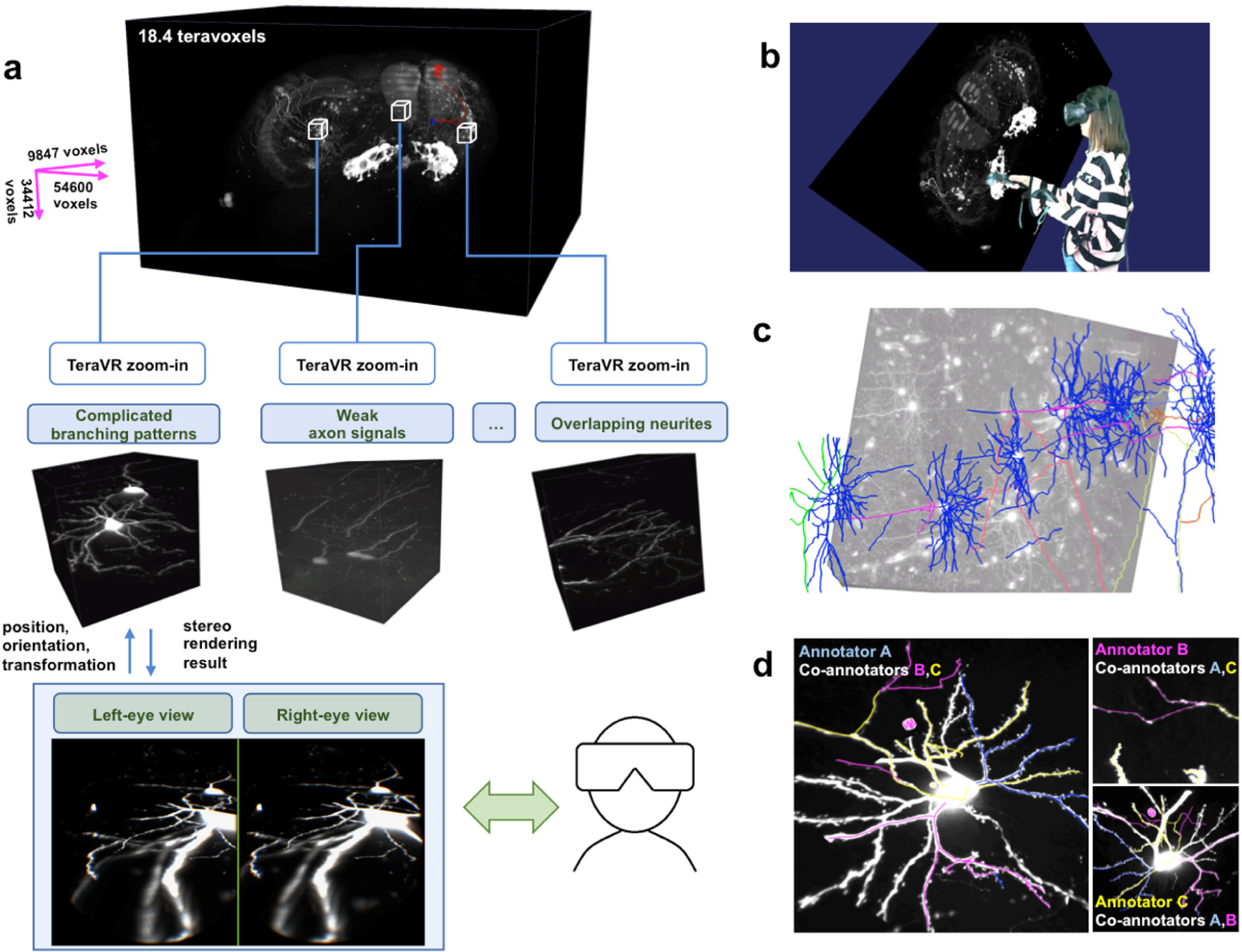
The overall scheme of TeraVR. (a) TeraVR is applicable to very challenging visualization and reconstruction scenarios such as complicated branching, weak signals, and overlapping neurites. With TeraVR, a user is able to combine stereoviews to observe the complex 3-D neurite patterns easily and perform the reconstruction effectively. Combining such visualization and data-exploration functions with terabyte-scale imaging data (e.g. whole-brain scale) management and streaming capability enables reconstruction of complex neuronal morphology at an unprecedented accuracy and efficiency. (b) A mixed reality visualization that demonstrates the use of TeraVR. Immersed in a virtual environment, the user manipulates the imaging data with TeraVR in a way similar to manipulating a physical object. (c) Multiple densely packed neurons from an image with high, noisy background intensity level were reconstructed using TeraVR. (d) Real-time collaboration is demonstrated by showing views from all participating annotators. Each annotator logs onto the cloud and adopts a unique color for both annotation and an avatar representing the user’s real-time location. The left figure shows the view for annotator A (blue), in which two avatars of co-annotator B (purple) and C (yellow) are seen. Annotation results are instantly shared among them. The upper right subpanel: annotator B examined a partially traced segment by co-annotator C, only to identify more branches after turning up the contrast and having a close-up view of the segment (without affecting the views of other annotators); bottom right subpanel: the view of annotator C.

To use *TeraVR*, a user wears a virtual reality headset (bottom right of **Fig. 1a**) and works within a virtual space defined for the brain image along with the neuron reconstruction and other location-references on the image. *TeraVR* generates synchronized real-time rendering streams for both left and right eyes (bottom left of **Fig. 1a**), which simulate how a person perceives real-world objects and thus forms stereo-vision. In this way, *TeraVR* facilitates efficient immersive observation and annotation (**Fig. 1b**, **Supplementary Video 12**) of very large-scale multi-dimensional imaging data, that can have multiple channels or from different imaging modalities (**Supplementary Fig. 1**). With the accurate pinpointing capability in *TeraVR* (**Supplementary Fig. 2**), in real-time a user can precisely and efficiently load the data of a desired high-resolution ROI to see detailed 3-D morphological structures (**Supplementary Fig. 2c**).

A user employs *TeraVR* to gain unambiguous understanding on a considerable number of challenging regions which typically contain complicated branching patterns, weak and discontinuous axon signals, overlapping neurites, etc. (middle of **Fig. 1a**) that are otherwise very hard, if not impossible, to distinguish confidently using any existing non-immersive visualization tools. *TeraVR* provides comprehensive tools for neuron reconstruction. In addition to single neurons, *TeraVR* was also used to reconstruct multiple densely packed neurons in very noisy images (**Fig. 1c**). *TeraVR* also allows multiple annotators working on the same dataset collaboratively using a cloud-based data server (**Fig. 1d**), in a way similar to Google-Docs, to combine multiple users’ input together efficiently.

We tested *TeraVR* in challenging situations for conventional non-VR approaches due to densely labeled and weakly imaged neurites. Such non-VR approaches include many visualization and annotation functions already existing in *Vaa3D* and *TeraFly*, as well as in other software packages such as ImageJ/Fiji (https://fiji.sc/) and Neurolucida (MBF Bioscience). First, for a strongly punctuated and highly intermingled axon cluster (**Fig. 2a**), five independent annotators reduced the time in tracing by 50-80% when they used *TeraVR* compared to *TeraFly*, the most efficient non-VR approach we found for these testing cases (**Fig. 2b**). Second, for exceedingly weak neurite signals (**Fig. 2c**), with *TeraVR* these annotators could consistently generate a neurite tract (bounded by branching points and/or terminal points) within 50 seconds, about 10 times better than the non-VR approach (**Fig. 2d**). For these weak signals, even when sometimes annotators needed to adjust the contrast in the visualization in both *TeraVR* and non-VR approaches, it was much easier for the annotators to use *TeraVR* than the non-VR method to find the right angle of observation and to add annotations on top of the signals. *TeraVR* reduced 60%∼80% of labor when measured with alternative metrics such as the number of strokes to complete a neurite tract in drawing (**Fig. 2d**). Third, for 109 dense or weak tracts, with *TeraVR* these annotators rarely needed more than 50 seconds to reconstruct any of such difficult tracts, while the non-VR approach normally needed about 10 times of effort for the same task (**Fig. 2e**). In 37.6% of tracts in this testing set, at least one annotator was not able to use the non-VR approach to reconstruct (**Fig. 2f**) while none of these annotators had trouble to accomplish the goal when *TeraVR* was used.

**Figure 2.**
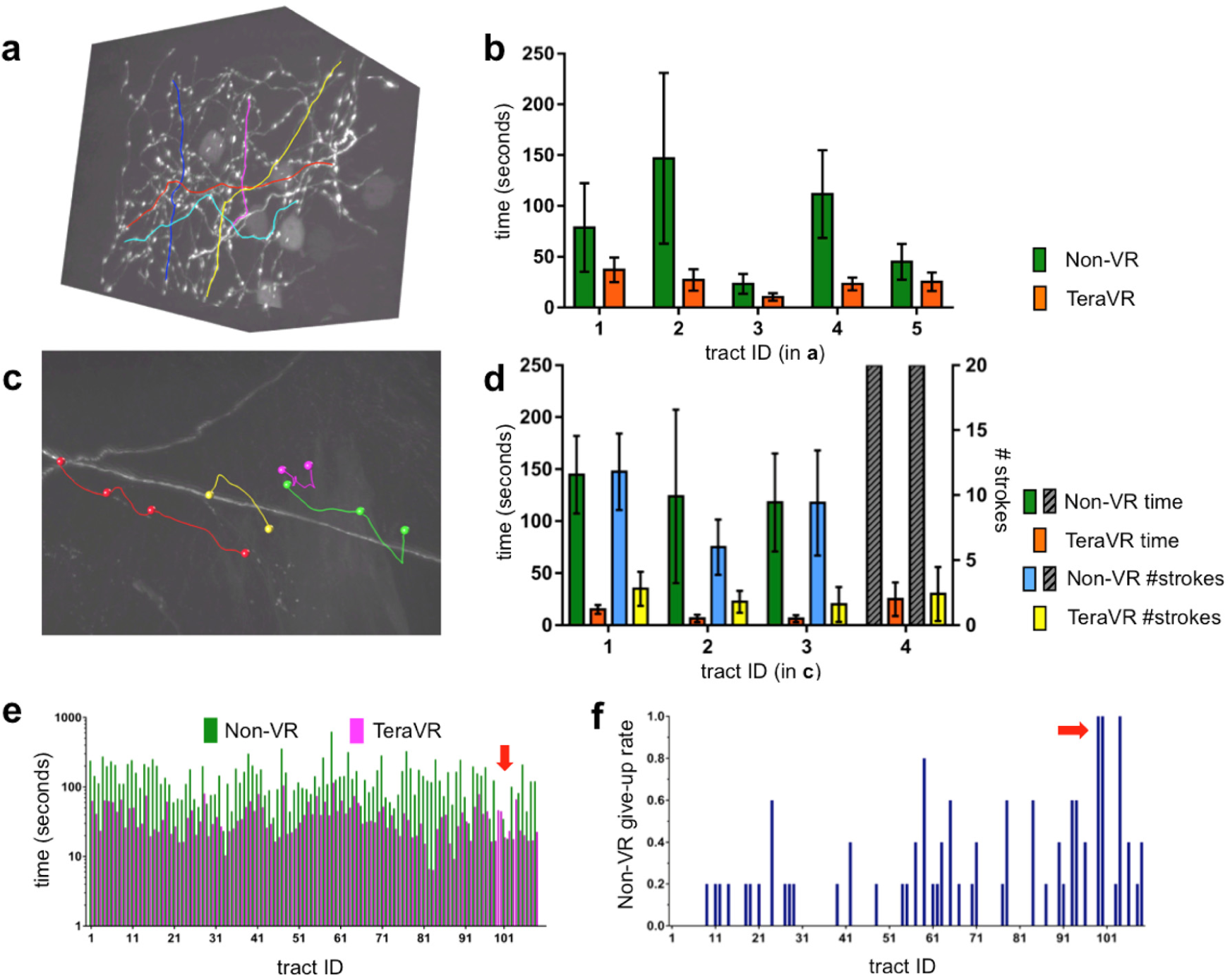
Efficiency of TeraVR. (a) A complex 3-D image volume with a number of intermingled, broken, strongly punctuated axon tracts. (b) Time spent to generate the five tracts in (a), each of which was produced by five independent annotators; the ‘non-VR’ results showed were obtained using TeraFly (same below in this figure); error bar: S.D. (c) A 3-D image volume with weak signal and strong noise, and the respective TeraVR reconstructions of barely visible neurite tracts. (d) Time and the number of operations needed to produce the tracts in (c). Gray bar: unavailable results (time/ number of strokes) for non-VR approach; error bar: S.D. (e) Average time of 5 annotators to generate 109 tracts that were hard to reconstruct. For non-VR, the average was calculated among the sub-group of annotators who succeeded in reconstructing the tract. (f) The give-up rate of non-VR for each tract in (e); an annotator was allowed to give up the attempt after trying 300 seconds; the give-up rate for each tract was defined as (#failed attempts)/ (#all attempts). Arrows in (e) and (f): the cases where no non-VR attempt was able to produce the respective neurite tracts.

A neuron may contain thousands or more neurite tracts, each of which is bounded by a pair of critical points, e.g. branching points, axonal or dendritic terminals, or the cell body (soma). Neurites are organized into local dendritic arbors, local axonal arbors, long projecting axon fibers, and distal axonal arbors. While some structures such as the major dendritic branches may be reconstructed using non-VR approaches, many other challenging cases (e.g. **Fig. 2**) will require the VR module in *TeraVR* for faithful and efficient reconstruction. Therefore, in *TeraVR* we designed a smooth switch between the VR mode and the non-VR mode to allow an annotator to choose a suitable mode to observe the imaging data and reconstruct neurites for different areas in a big imaging dataset.

This technology allowed us to reconstruct complete 3-D morphology of neurons from the whole mouse brain, each of which was repeatedly curated by four to five annotators to ensure accuracy (**Figs. 3** and **4**, **Supplementary Fig. 3 and 4)**. To better understand the usability of *TeraVR*, we trained 15 annotators to independently produce complete reconstructions for different types of neurons. We analyzed under which situations these annotators would switch between VR and non-VR modes to understand the strength of the VR-mode (**Fig. 3**). VR was used mostly in densely arbored areas such as axonal arbors and sometimes also in local dendrites (**Fig. 3a**, and **Supplementary Figs. 3a ∼ 3c**). The areas done by VR often have low or very low signal-to-noise-ratio (SNR) (**Fig. 3a, Supplementary Fig. 3**, **Methods**). For 44 thalamic neurons in two mouse brains, the percentage of very low SNR regions correlated linearly with the VR-portion of neurons (**Fig. 3b**). Linear correlation was also observed in analyzing 73 neurons in caudate putamen in the two brains (**Fig. 3c**). For all these 117 neurons together, over 90% of VR usage was dedicated to the reconstruction of neurites in the below average SNR regions (**Fig. 3d**).

**Figure 3.**
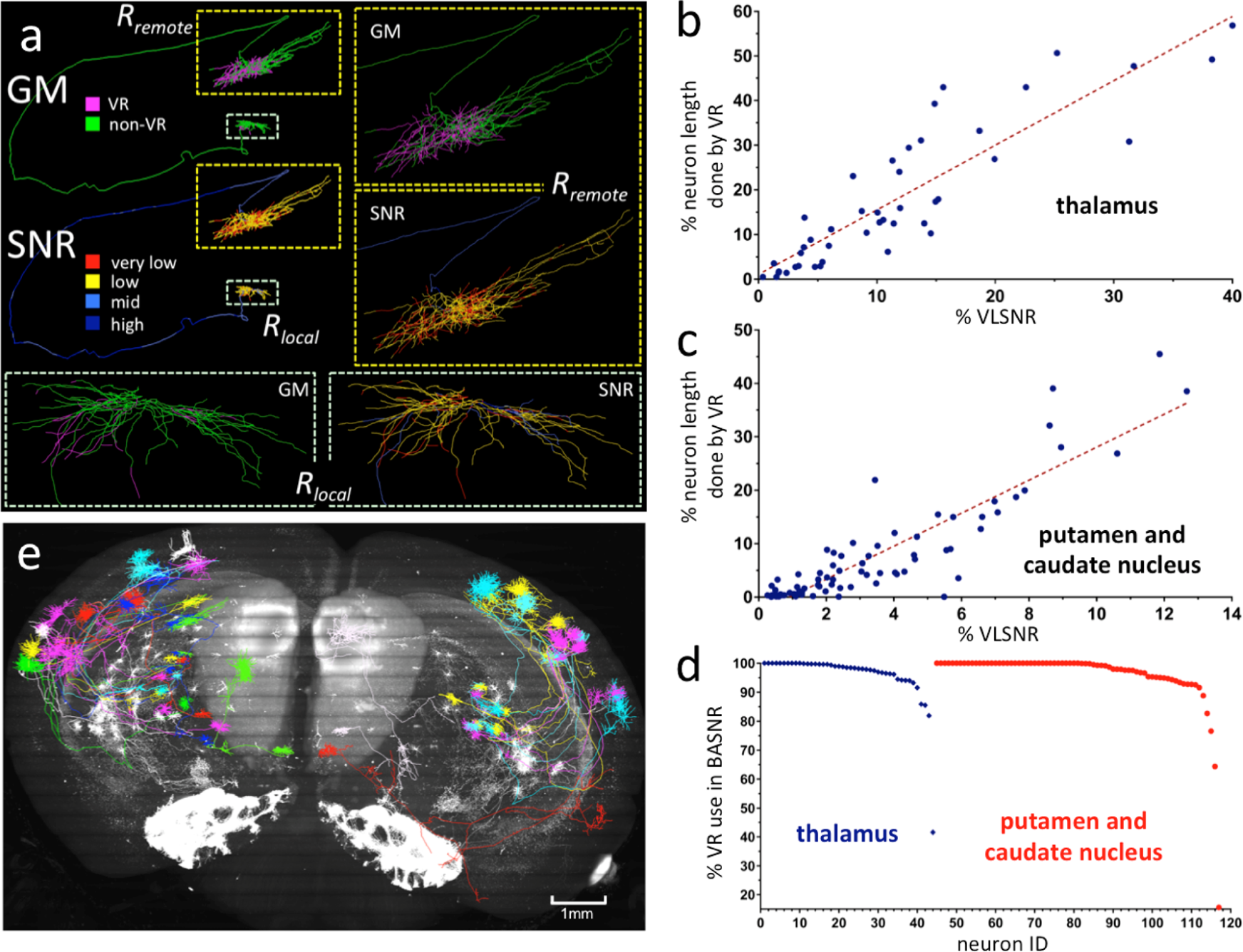
Complete reconstruction of neurons at whole brain scale using TeraVR. (a) A thalamic cell reconstructed using TeraVR. Upper left: a complete reconstruction of the neuron color-coded using ‘GM’ (Generation Method) and ‘SNR’ (signal-to-noise-ratio) schemes; in ‘GM’, magenta and green colors stand for neurites reconstructed using VR and non-VR, respectively; in ‘SNR’, blue, sky blue, yellow, and red colors indicate neurites with high, mid, low and very low SNR, respectively; two close-up views of local dendrites and remote axons are also shown in the right and the bottom. (b) For a set of 44 completely reconstructed thalamic neurons (33 from brain No. 17302, 11 from brain No. 17545), the correlation between the portion of a neuron traced using the VR mode of TeraVR and the portion of this neuron that has very low SNR (VLSNR). (c) For a set of 73 completely reconstructed neurons in caudate putamen (58 from brain No. 17302 and 15 from brain No. 17545), the correlation between the portion of a neuron traced using the VR mode of TeraVR and the portion of this neuron that has very low SNR (VLSNR). (d) The use of VR mode in reconstruction of BASNR (below average SNR) regions in each of the 117 neurons. (e) Whole-brain plot of 33 thalamic neurons reconstructed from brain No. 17302; gray: maximal intensity projection of this brain image; color-code: each neuron in a randomly assigned color.

**Figure 4.**
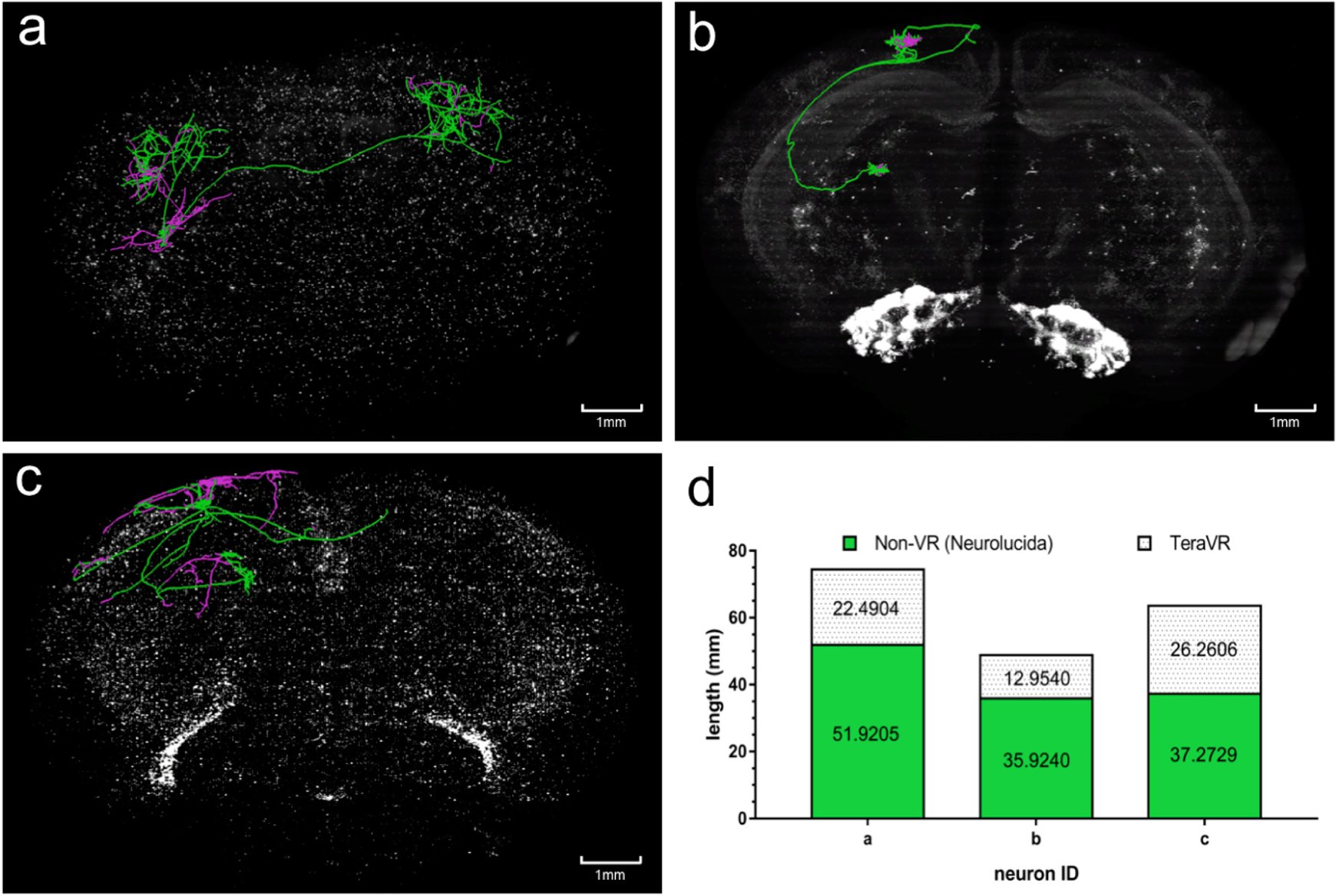
The use of TeraVR in validating, correcting, and extending complex neuron-reconstructions produced with Neurolucida. (a) ∼ (c) Three examples of reconstructed neurons overlaid on the whole-mouse brain imaging data, from three different brains (IDs: 236174, 17545, 17300), respectively. Green: initial reconstructions produced using Neurolucida; magenta: recovered missing portion of reconstructions using TeraVR. (d) The length of neuron reconstructions produced for (a)∼(c), respectively.

We further investigated whether reconstructions of similar accuracy could have been produced using other commonly used tools. We used *TeraVR* to recheck the reconstruction of neurons with very complex morphology, such as the cortico-cortical neurons, initially generated by annotators who had a lot of experience in using a popular reconstruction tool called *Neurolucida* (*Neurolucida 360 or NL360)*. Since *NL360* does not have comparable capability to handle big data IO streaming, the annotators needed to load a portion of the imaging data at a time to reconstruct neurons, at a much slower pace. More importantly, upon rechecking in *TeraVR* we found imperfectness of these *NL360*-based reconstructions (**Fig. 4 a-c**). The under-tracing of missing neurites was most notable, and the topology errors and over-tracing were common (**Fig. 4 a-d**) even for the cells traced from overall clearly labeled brains. In some cases, more than 40% of neurites of a neuron were found to be missing (**Fig. 4 c-d, Supplementary Fig. 4a**). Notably, it was often seen a missing axonal branch at the proximal part of an axon, which indicated missing a long projection and the corresponding whole distal targeting axonal cluster (**Fig. 4a, Fig. 4c**). Also, annotators could choose to proceed along a wrong direction when a confusing branching region was encountered, which would lead to more severe reconstruction errors (**Supplementary Fig. 4**). These indicate the limitation of conventional tools for accurately observing neuronal structures in certain special situations such as dense neurites, axonal collaterals in dendrosomatic regions, where signals become obscure (for example, long axonal collaterals extending along pia, **Fig. 4c**). This observed limitation is common for the non-VR approaches, such as *Vaa3D-TeraFly* and *Neurolucida*, compared to *TeraVR*. A careful examination of 17 complex neurons from three whole-brains indicated that *TeraVR* extended 10-103% of the overall lengths of reconstructions from these neurons (**Supplementary Table 1**). We also carefully examined several other VR software packages and did not find any one that had comparable functions as *TeraVR* (**Supplementary Tables 2 and 3**).

In contrast to 2-D display devices in front of which multiple people may view the same visualization simultaneously, currently one 3-D VR headset can only be worn by one person at a time, therefore an annotator may not communicate easily with others once this person is working in the VR environment. To overcome this limitation, in *TeraVR* we developed a collaboration mode with which multiple users can join the same session to reconstruct the same neuron at the same time, similar to the co-editing feature of Google-Docs. Specifically, in *TeraVR* we implemented a cloud-based server-client infrastructure, with which the annotation data of individual annotators are streamed to the server in real time and merged with the data produced by other collaborating annotators. Users are able to see all annotations produced by others in real time and perform certain further annotations. We assembled a geographically remote team of annotators in Nanjing (China), Shanghai (China), and Seattle (USA) to use this collaboration mode to simultaneously reconstruct complicated 3-D neuron morphology from the whole-brain imaging dataset (**Fig. 1d** and **Fig. 5**). Three annotators, each from a different city, were able to co-reconstruct in real-time dendritic and axonal structures around the soma of a neuron (**Fig. 5 a-c**) with only 20% of time compared to one single annotator (**Fig. 5e**). A Sholl analysis (Langhammer, et al, 2010) indicated the *TeraVR*-reconstructions produced by different combinations of annotators had consistent topology (**Fig. 5d**). A length analysis indicated the difference of neuron-lengths generated by such combinations of annotators was also small, at only 0.77% of the average total length of the reconstruction (**Fig. 5e**). A spatial distance analysis indicated the average lateral apartness of these reconstructions was about 3.5 voxels, which was 0.05% of the longitudinal span of the neuron (**Fig. 5f**). This study indicates the power of *TeraVR*’s collaborative approach for remote annotation.

**Figure 5.**
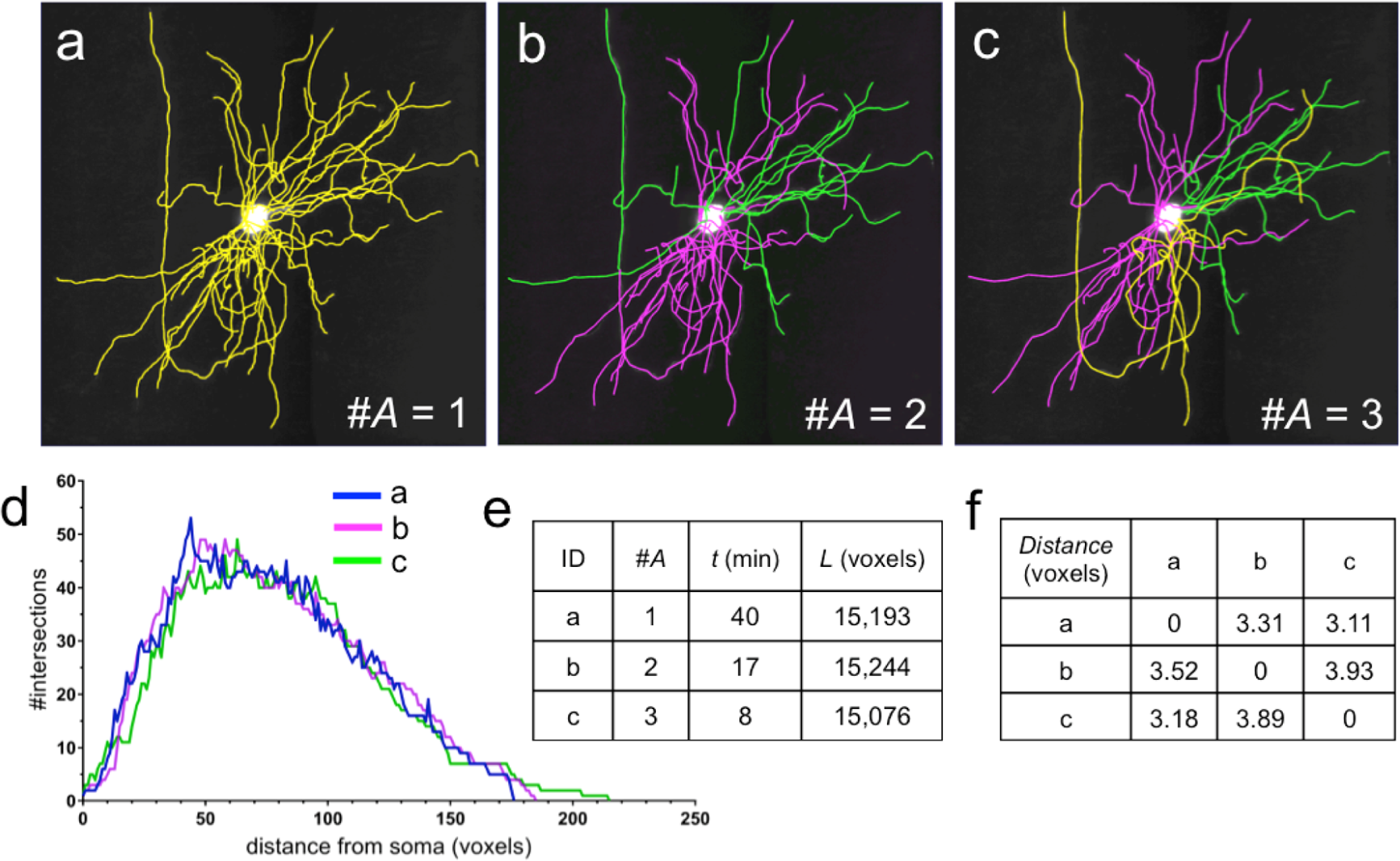
Results produced using the collaboration mode of TeraVR. (a) ∼ (c) Reconstructions done by different numbers of collaborating annotators; different colors of neurites indicate parts done by different annotators; #A: number of annotators. (d) Sholl analysis of three reconstructions in (a)∼(c). (e) Summary of the number of annotators, reconstruction time, and the total length of reconstructions in (a)∼(c). (f) The pairwise spatial distance of reconstructions in (a)∼(c).

We developed *TeraVR* as an open system, which can be augmented by a number of other programs without compromising its modularity. In particular, we enhanced *TeraVR* using several artificial intelligence techniques to further improve the efficiency of annotators. First, for the imaging data, we trained a deep-learning model, U-Net (Ronneberger, et al 2015), based on high-quality reconstructions produced using *TeraVR*; then in *TeraVR* we allowed a user to quickly invoke the trained U-Net to separate neurite signal from background (**Supplementary Figs. 5a, 5b**). We streamed the U-Net filtered images in real-time to *TeraVR* as an option that a user could choose. This U-Net model could also be iteratively refined based on user’s feedback, thus it could be adapted to different brain images when needed. Second, for neuron reconstructions, in *TeraVR* we implemented a data-filtering model to detect various outlier structures, such as branches that had sharp turns (e.g. turning angle greater than 90 degrees or 135 degrees or other user-specified values), and then generate alerts to allow users to immediately focus on the structures that might be traced with errors (**Supplementary Figs. 5c, 5d**).

## Discussion

*TeraVR* offers an immersive, intuitive and realistic experience for exploring brain imaging data, similar to the mixed reality visualization shown in **Fig. 1b** and **Supplementary Video 12**, where real and virtual contents were synthetically put together to demonstrate the user experience of *TeraVR*. While VR has not been widely used in biology, it is useful for biological problems especially due to the intrinsic multidimensional nature of many biological datasets, and has the potential to be integrated as the next standard protocol. *TeraVR* is among the first demonstration of such utility with great potential. While immersive VR visualization of biological surface objects and sometimes also imaging data were shown in applications such as biological education and data analyses (**Supplementary Table 2**), there is little existing work on developing open-source VR software packages for very complicated and teravoxel-scale imaging datasets such as the whole brain imagery as we have introduced here. We expect that *TeraVR* can also be used to analyze other massive-scale datasets especially those produced using fast or high-resolution microscopy methods, such as the light-sheet microscopy (Keller, et al, 2008; Ahrens, et al, 2013; Silvestri, et al, 2013), expansion-microscopy (Chen, et al, 2015), and recent nanoscale lattice microscopy (Gao, et al, 2019).

We chose to focus on applying *TeraVR* to the whole-brain single-neuron reconstruction challenge for two major reasons. First, currently no other alternative tools are able to reconstruct the fine, distal arborizations of neurons unambiguously in this way. Second, there has been little previous work on streamlining the large-scale data production of the complete single-neuron morphology at high precision and also at whole-brain scale. *TeraVR* has been a crucial tool to help several teams reconstruct precisely hundreds of full morphologies, with various image qualities, not only for single neurons but also for multiple densely packed neurons in very noisy images. These reconstructions have been released to the public databases e.g. NeuroMorpho.Org and the BRAIN Initiative Cell Census Network initiative.

Two additional aspects of *TeraVR* make this software package unique: the collaboration mode and the integration of the artificial intelligence methods. *TeraVR* users can readily work together remotely and curate each other’s reconstructions. Such real-time ensemble-annotation greatly improves the consistency, robustness, accuracy, speed, and actual fun of neuron reconstruction. With the further help of machine learning-based data analysis modules in both image and reconstruction domains, *TeraVR* will allow effective crowdsourcing and production of large-scale “gold standard” reconstructions, which in addition to its inherent value will further help the automation of neuron reconstruction and systematic studies of neuron morphometry.

## Acknowledgement

H.P. conceived, initialized and managed this project, co-developed the design of the software and wrote the paper. Y.W. developed the *TeraVR* system, conducted the experiments and data analysis with helps from the team, and also together with H.P. wrote the paper. Q.L. and L.K. helped developing the *TeraVR* system. L.L., Yun W. and Z.Z. helped on the data production in this manuscript. N.Z., R.C., X.L., Y.G., M.H., Z.G., and W.X. supported the project and offered resources to accomplish this research. Qingming L. collaborated with H.Z. to provide whole mouse brain imaging data. We thank Wenbin Wang, Yufei Jin, Peng Wang, Shengdian Jiang, Qiang Ouyang, Sujun Zhao, Yuanyuan Song, Lulu Yin, Jia Yuan, Guodong Hong, Wan Wan, Xuefeng Liu, Linus Manubens-Gil, Peng Xie, Hsienchi Kuo, Zongcai Ruan, Alessandro Bria, and many other members in the collaborating organizations for a great amount of support of data, algorithm, testing and feedback.

## Methods

### Data preparation

*Tnnt1-IRES2-CreERT2;Ai82;Ai140* (brain ID No. 17302 and 17545), *Gnb4-IRES2-CreERT2;Ai139* (No. 236174) and *Plxnd1-CreER;Ai82;Ai140* (No. 17300) mice were used in fMOST (Li, et al, 2010) imaging to produce raw image stacks, which were further converted into the *TeraFlY*-format using Vaa3D’s module TeraConverter.

### *TeraVR* visualization

*TeraVR* provides an immersive VR environment and true 3-D experience for interactive neuronal image visualization and annotation. A VR device, e.g. the HTC Vive (https://www.vive.com/us/), typically has a wearable headset (also known as head-mounted device) with two independent monitors. The left monitor is exclusively viewed by the left eye; and the right monitor by the right eye. *TeraVR* produces and feeds two slightly different rendering streams for left and right monitors, which are viewed by the user simultaneously to create a realistic stereo visualization. We used the ray-casting technique to render neuronal volume images. To allow the user to observe the data inside of the image volume, *TeraVR* adds a clipping plane orthogonal to the view direction to the typically used cube-model texture mapping to form a closed surface.

### Collaboration Mode

*TeraVR* allows multiple annotators to work collaboratively during reconstruction. To enable the collaboration mode, a collaboration server is deployed on the cloud or in the intranet. The server receives messages from each connected annotator, and broadcasts to the others. An annotator joins a collaboration session by specifying the username and the IP/port of the collaboration server. Once connected, the annotator’s real-time working location in an image will be represented by an avatar, which is visible to all the other annotators. The annotator is also assigned a unique color, which is used as both the avatar’s color and the annotation’s color. When the annotator edits the reconstruction, e.g. adding / deleting a neurite or a marker, the operation is converted to a globally understandable command, which is sent to the server. The server maintains a queue of commands and dispatches them in sequence to all the connected annotators. In this way, the reconstruction result is synchronized among all the annotators.

### Mixed reality video making

To generate a mixed reality demonstration (**Fig. 1b**) that shows how *TeraVR* works, we first setup a physical camera to capture the movement of the annotator. A green screen was used to help remove the background. Meanwhile, an additional virtual camera was placed at the location of the physical camera (rather than being mounted on the VR headset) to generate a rendering stream of *TeraVR* from a third-person view. Importantly, the physical and the virtual cameras had exactly the same settings, including position, orientation, focus, etc., so that the real video stream was directly superimposed over the virtual one. These two cameras were started after *TeraVR* was launched. The mixed reality video was produced by synthesizing these two video streams.

### Profiling the image quality of a neuron

To evaluate how hard to reconstruct a neuron, we profiled the underlying image quality for a neuron. We first decomposed a neuron structure into a set of segments, each being bounded by a pairs of critical points (branch points, terminal points, or the soma). The foreground (***F***), background (***B***), and critical background (***B_crt_***) were extracted for each segment: ***F*** was defined as the area enclosed within the radius of reconstructed neurite segment, ***B*** was defined as the bounding box of the segment excluding ***F***, and ***B_crt_*** was defined the 20% brightest voxels within ***B***. We then calculated the signal-to-noise-ratio (**SNR**) for a neurite segment as 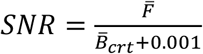, where 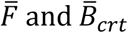 were the average intensities for the image-voxels in foreground and critical background, respectively. Four SNR ranges were defined based on annotators’ consensus opinions: “very low” for SNR ∈ (−∞, 1.0] (the neurite signal was either very weak or very noisy), “low” for SNR ∈ (1.0, 1.2] (the signal was still in low quality), “mid” for SNR ∈ (1.2, 1.4], and “high” for SNR values ∈ (1.4, ∞) (strong signals, which are clear and easy to trace). The overall image SNR of a neuron was calculated as the segment-wise average SNR weighted by the length of each segment.

### Computer configuration

*TeraVR* was implemented and evaluated on computers with Intel Core i7-7700 CPU @ 3.60GHz, 64GB memory, NVIDIA GeForce GTX 1070 GPU, Windows 10 64-bit edition, and uses HTC Vive as the virtual reality device.

### Compatibility

*TeraVR* can be used to explore multi-dimensional, multi-channel image data, as long as the data format is supported by Vaa3D. For very large-scale images (>100 billion voxels), it is recommended to organize the data in the Vaa3D-Terafly format for smooth performance.

### Availability

*TeraVR* is released Open Source, as part of Vaa3D. A user guide for *TeraVR* is provided in **Supplementary Note 1**. Whole-brain test data is available upon request due to their very large sizes.

---

**Supplementary Figure 1.**
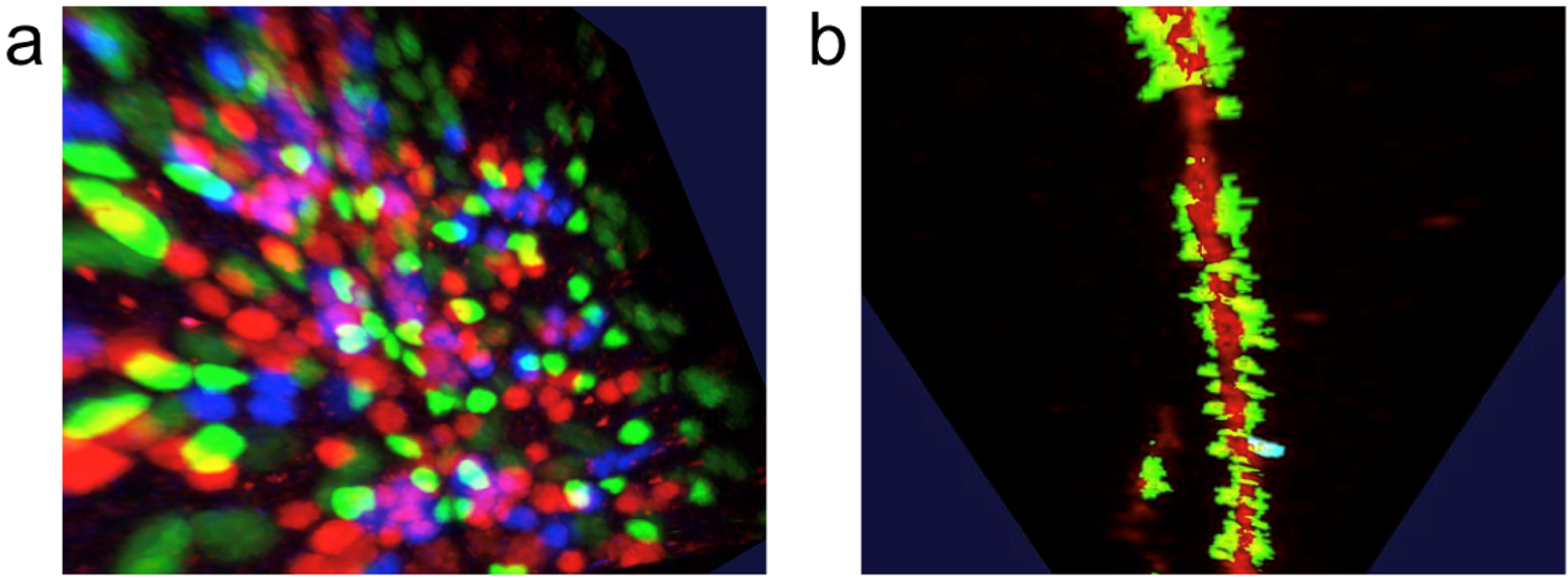
Visualization of images of multiple channels using TeraVR. (a) A group of cells labeled in red, green, and blue colors. (b) An image stack in which the dendritic neurites are visualized in red color, and the spines are visualized in green color.

**Supplementary Figure 2.**
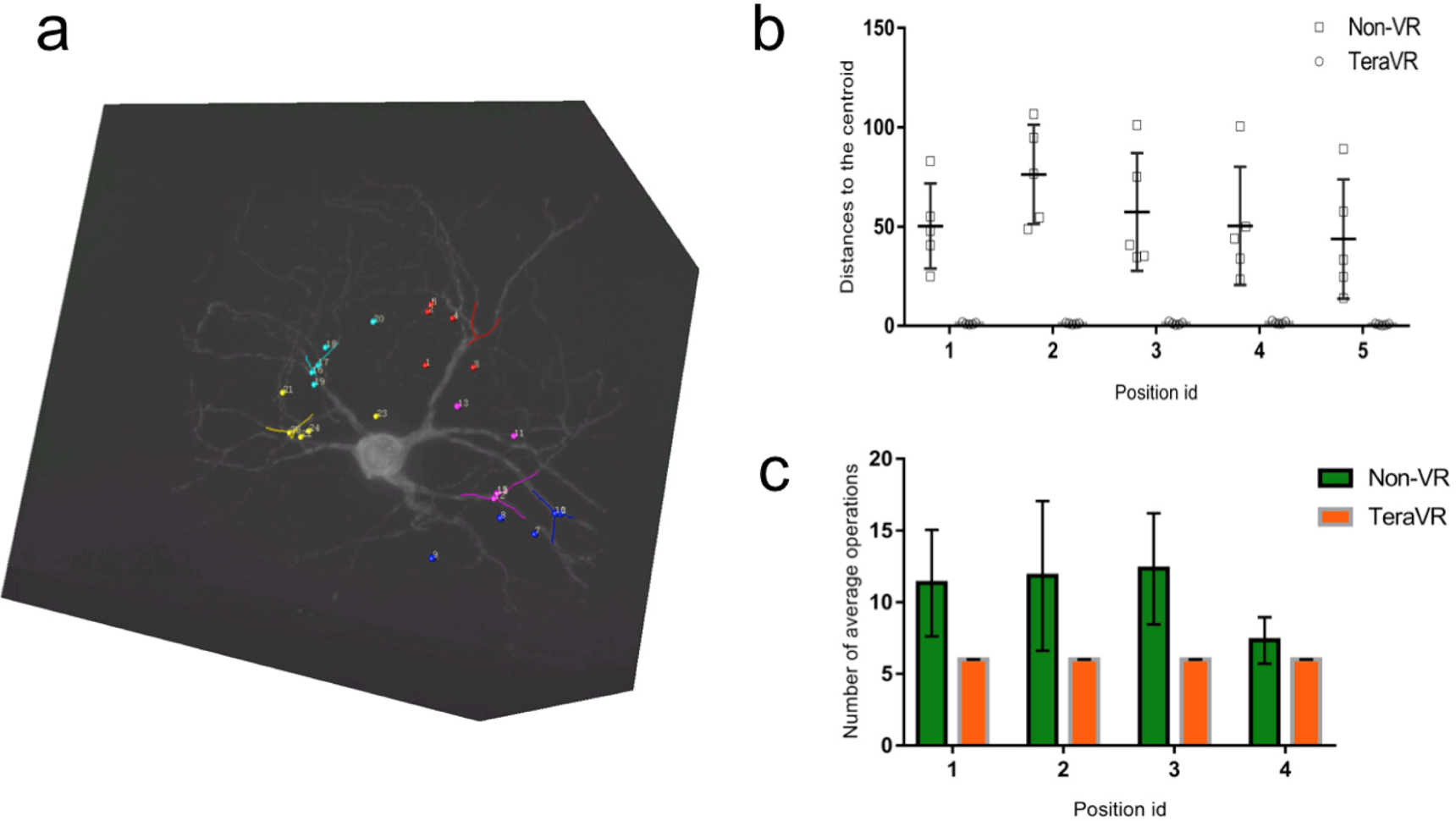
Accurate pinpointing in TeraVR. (a) A local image volume that contains a dendritic tree. 5 bifurcations are highlighted using Y-shaped structures of different colors. 5 attempts for adding a marker at each of the bifurcations are made using TeraVR and non-VR approach, respectively. The displayed markers are the according attempts to pick up the bifurcations using non-VR approach. (b) For each group of attempts, a geometric centroid is calculated. The plot shows the distance to the centroid; error bar: S.D. (c) A plot of number of operations needed in order to go to the highest resolution of a ROI from the lowest resolution. TeraVR has stable performance and requires only the fewest number of operations. The non-VR approach lacks enough accuracy for pinpointing and thus needs more operation to accomplish the task; error bar: S.D.

**Supplementary Figure 3.**
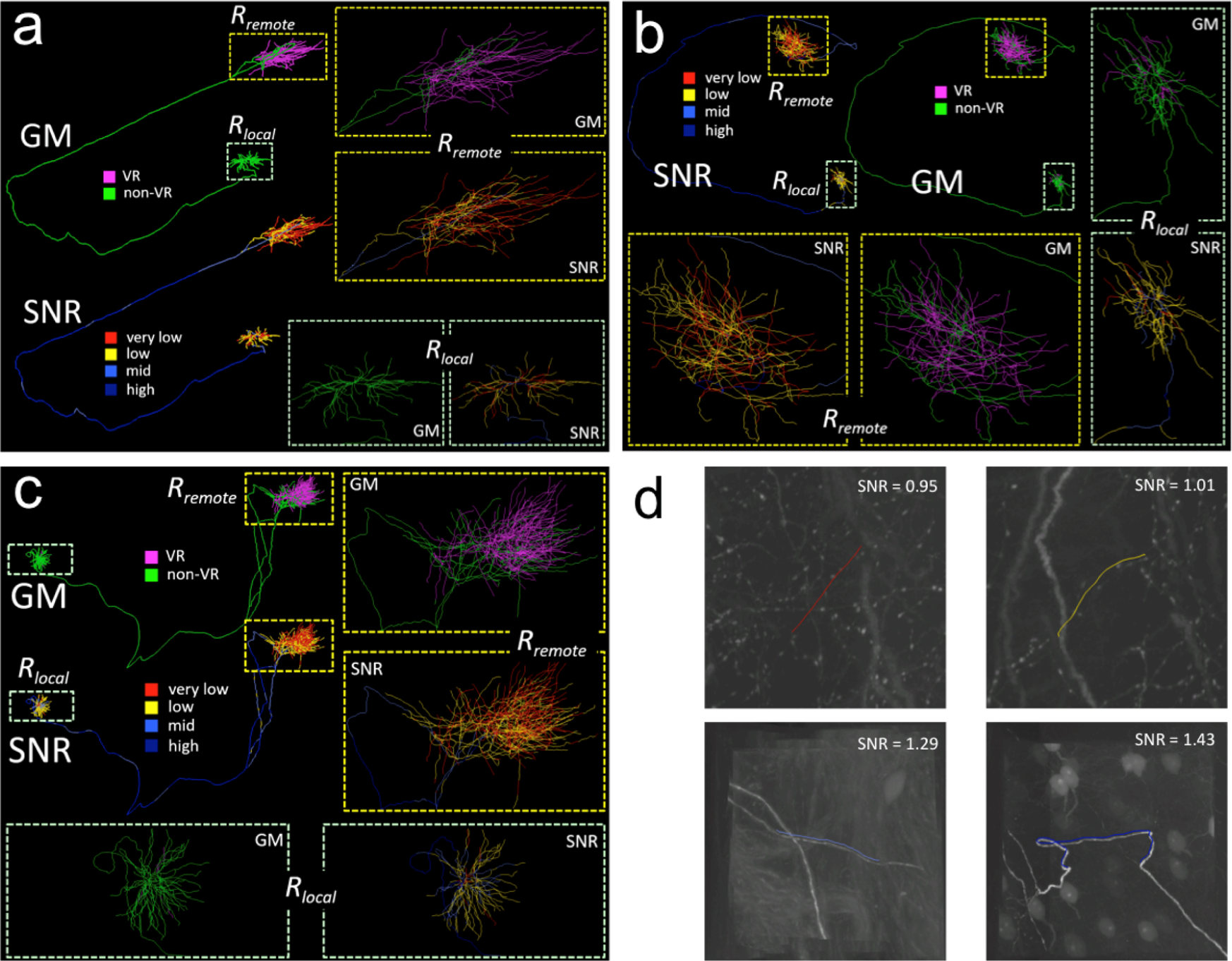
More complete reconstructions of neurons at whole brain scale using TeraVR. (a)-(c) 3 more neurons reconstructed using TeraVR. Refer to Figure 3 for the meaning of the color-coding. (d) The illustrations of image regions with various SNRs. The neurites are given a slight offset for clarity.

**Supplementary Figure 4.**
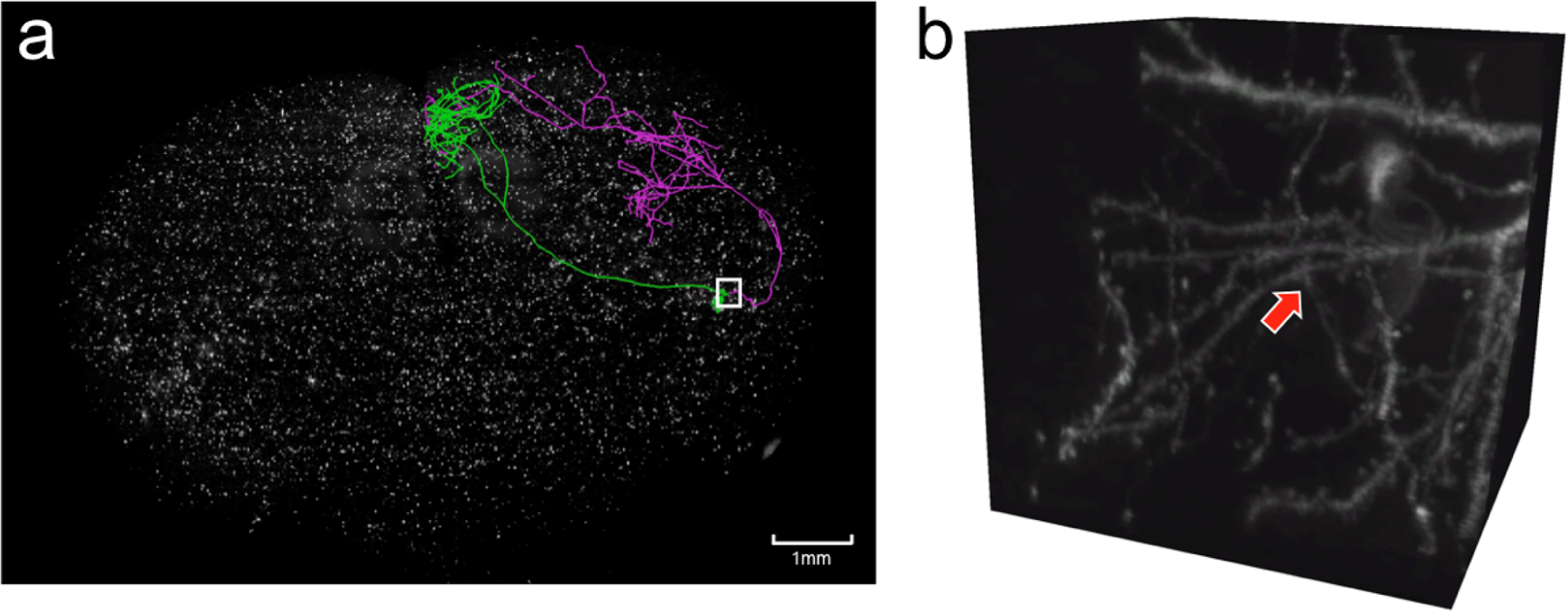
TeraVR helps picking up a major missing cluster in neuron-reconstruction produced first with Neurolucida. (a) A neuron first produced with Neurolucida (green) and then corrected by TeraVR (magenta), overlaid on the whole-mouse brain imaging data. The image quality is challenging in the region specified by the white box, leading to a large size of incorrectly-traced arbor in the Neurolucida reconstruction that was later identified and deleted using TeraVR (the deleted part of the Neurolucida reconstruction is not displayed here). (b) Full-resolution imaging data corresponding the to the white box in (a). The red arrow points to the critical position that corresponds to the reconstruction error resulting in a major missing cluster.

**Supplementary Figure 5.**
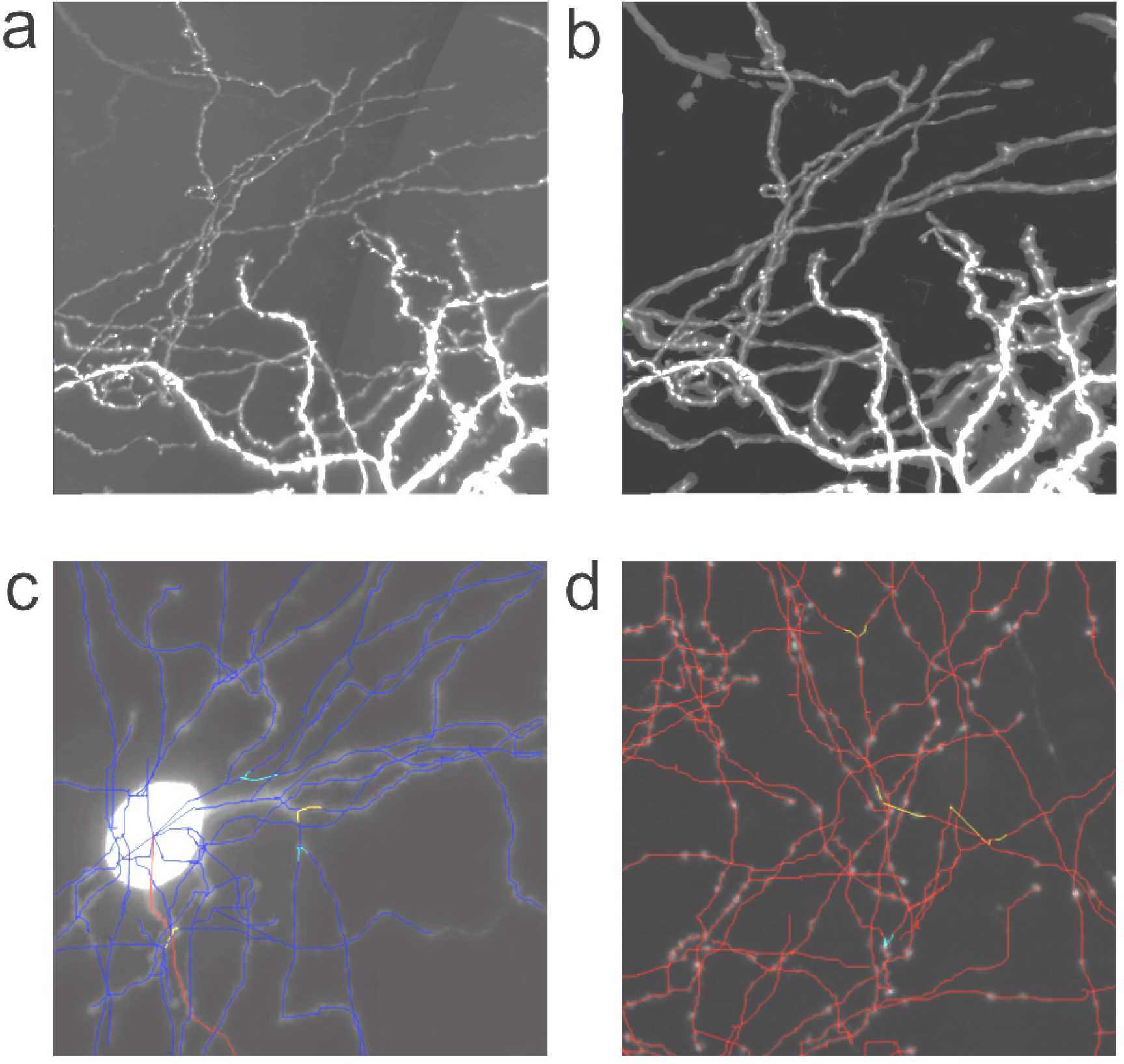
Enhancing TeraVR using several AI modules. (a) The original image visualized in TeraVR. (b) The U-Net optimized image visualized in TeraVR. (c) A partial dendritic tree, where bifurcations with abnormal angles are highlighted (non-blue/red colors). (d) A terminal axon arbor, where bifurcations with abnormal angles are highlighted (non-red colors). For all subfigures, brightness +40%, contrast −40% for more visibility.

**Supplementary Table 1.** Statistics for 17 neuron-reconstructions (from three brains) that were first reconstructed using Neurolucida and then corrected using TeraVR.

| Brain ID | Neuron ID | Total length (mm) | TeraVR length (mm) | Non-VR length (mm) | % increased |
| --- | --- | --- | --- | --- | --- |
| 236174 | 04229-04328-X13663-Y8589 | 67.6354 | 34.3252 | 33.3102 | 103.0471% |
| 17300 | 5969-X27278-Y20820 | 63.5335 | 26.2606 | 37.2729 | 70.4549% |
| 17300 | 03514-3525-X19676-Y45282 | 51.6424 | 16.1144 | 35.528 | 45.3569% |
| 236174 | 03329-03428-X13938-Y26099 | 74.4109 | 22.4904 | 51.9205 | 43.3170% |
| 17545 | 05574-X24399-Y33944 | 48.878 | 12.954 | 35.924 | 36.0595% |
| 236174 | 03536-03545-X15159-Y25525 | 93.6728 | 19.3034 | 74.3693 | 25.9561% |
| 17545 | 05689-X21900-Y16152 | 21.3688 | 3.85913 | 17.5096 | 22.0401% |
| 17545 | 06151-X24259-Y36270 | 21.5348 | 3.85515 | 17.6797 | 21.8055% |
| 236174 | 03529-03628-X12805-Y10541 | 106.858 | 18.2958 | 88.5619 | 20.6588% |
| 236174 | 03001-03008-X12887 Y24248 | 115.714 | 18.3905 | 97.3237 | 18.8962% |
| 17545 | 06070-X20183-Y17777 | 27.1727 | 4.217 | 22.9557 | 18.3702% |
| 17545 | 06034-X23713-Y35681 | 42.3159 | 6.34982 | 35.966 | 17.6551% |
| 17545 | 05534-X20427-Y33851 | 54.3758 | 7.94209 | 46.4337 | 17.1042% |
| 17545 | 05996-X19743-Y18066 | 30.3852 | 4.07729 | 26.3079 | 15.4983% |
| 17300 | 03426-X20339-Y44872 | 65.7337 | 8.45904 | 57.2747 | 14.7692% |
| 236174 | 03429-03528-X12632-Y10625 | 97.0815 | 11.2848 | 85.7968 | 13.1529% |
| 236174 | 03447-03459 X12562 Y10626 | 73.5985 | 7.13707 | 66.4614 | 10.7387% |

**Supplementary Table 2.** VR software for scientific visualization. ^1^ syGlass has preliminary support for neuron reconstruction by placing consecutive nodes in the space to represent neurites, which is not practical and efficient for use in complete neuron reconstruction from whole-brain data. Ref_1: https://www.arivis.com/en/imaging-science/arivis-inviewr. Ref_2: https://www.syglass.io/. Ref_3: Stefani, C., Lacy-Hulbert, A., & Skillman, T. (2018). ConfocalVR: Immersive Visualization for Confocal Microscopy. Journal of Molecular Biology, 430(21), 4028–4035.

|  | arivis<br>InViewR | syGlass | ConfocalVR | VRNT | *TeraVR* |
| --- | --- | --- | --- | --- | --- |
| VR visualization<br>of 3D images | Yes | Yes | Yes | Yes | Yes |
| VR neuron<br>reconstruction | No | Preliminary <sup>1</sup> | No | Yes | Yes |
| Reported<br>application to<br>whole brains | No | No | No | No | Yes |
| License | Commercial | Commercial | Free only for<br>nonprofit | Free | Free and Open<br>Source |
| Reference/link | Ref_1 | Ref_2 | Ref_3 | [24] | This<br>manuscript |

**Supplementary Table 3.** A comparison between TeraVR and VRNT regarding the data management and compatibility, functions for reconstruction, and visualization and navigation. **Bold**: critical features.

|  | Features | VRNT | *TeraVR* |
| --- | --- | --- | --- |
| Data management and compatibility | Support for imaging data | Yes | Yes |
|  | **Size of tested imaging data (voxels)** | 300 megavoxels | 10~30 teravoxels |
|  | Support for multi-channel imaging data | No | Yes |
|  | Direct opening of morphology data | No | Yes |
| Functions for reconstruction | Adding or deleting tracts | Yes | Yes |
|  | Modifying the fine geometry of a tract | No | Yes. A number of functions such as subdivision and dragging are provided. |
|  | Undo | Yes | Yes |
|  | Redo | No | Yes |
|  | **Semi-automatic reconstruction** | No. Reconstruction results might not well align with the signal. | Yes. Virtual finger makes neurites align with signals. |
|  | **Collaborative reconstruction** | No | Yes |
|  | **Support for artificial intelligence** | No | Yes |
| Visualization and navigation | **Visualization mode** | Single. VR mode only. | Dual. Allow convenient switch between VR and non-VR mode. |
|  | **Multiresolution display** | No. Thus not applicable for whole-brain applications w/o redevelopment. | Yes. It is helpful when working on whole-brain imaging data. |
|  | **Good visibility of complex, weak and/or noisy regions** | No. Visibility is not good in many challenging cases. | Yes |
|  | Contrast adjustment | No | Yes |
|  | Hide reconstructions | No | Yes |
|  | Translating | Yes | Yes |
|  | Rotation | No | Yes |
|  | Scaling | No | Yes. It could be useful for anisotropic imaging data. |

